# A minimal tooth enhancer regulates *dlx2b* expression during zebrafish tooth formation: insights into *cis*-regulatory logic in organogenesis

**DOI:** 10.1101/2022.01.20.477116

**Authors:** William R. Jackman, Yujin Moon, Carol K. Cox, Drew R. Anderson, Audrey A. DeFusco, Vy M. Nguyen, Sarah Y. Liu, Elisabeth H. Carter, Hana E. Littleford, Elizabeth K. Richards, Andrea L. Jowdry, Yann Gibert

**Affiliations:** Biology Department, Bowdoin College, Brunswick, ME 04011, USA; Department of Cell and Molecular Biology, Cancer Center and Research Institute, University of Mississippi Medical Center, Jackson, MS 39216, USA

**Author notes:** Correspondence: William R. Jackman or Yann Gibert. Current address: Department of Medical and Molecular Genetics, University of Indiana, Indianapolis, IN 46202, USA.

**Keywords:** dlx2b, cis-regulation, tooth

## Abstract

Despite growing recognition of the importance of *cis*-regulatory elements in vertebrate development, the mechanisms by which enhancers control gene expression during organogenesis remain incompletely understood. To address this gap, we investigated the regulation of the transcription factor *dlx2b* during zebrafish larval tooth formation. Using CRISPR/Cas9-mediated genome editing, we generated a GFP knock-in line that recapitulates *dlx2b* expression in developing tooth germs. Through targeted manipulation of enhancer sequences, we identified a minimal tooth enhancer (MTE), which is sufficient to drive most of the endogenous *dlx2b* tooth germ expression pattern *in vivo*. Functional dissection of the MTE revealed that four evolutionarily conserved transcription factor binding sites are essential for enhancer activity. Mutating these sites within a transgenic reporter abolishes enhancer-driven expression, while deletion of the same sequences at the endogenous *dlx2b* locus causes a dramatic shift in the gene’s expression pattern. These findings suggest that loss of MTE function permits alternative *cis*-regulatory elements to gain control of the promoter, highlighting the dynamic nature of enhancer-promoter interactions during development. Together, these results uncover fundamental principles of enhancer function during vertebrate organogenesis and demonstrate the power of empirical dissection in decoding *cis*-regulatory architecture.

**Graphical Abstract:** 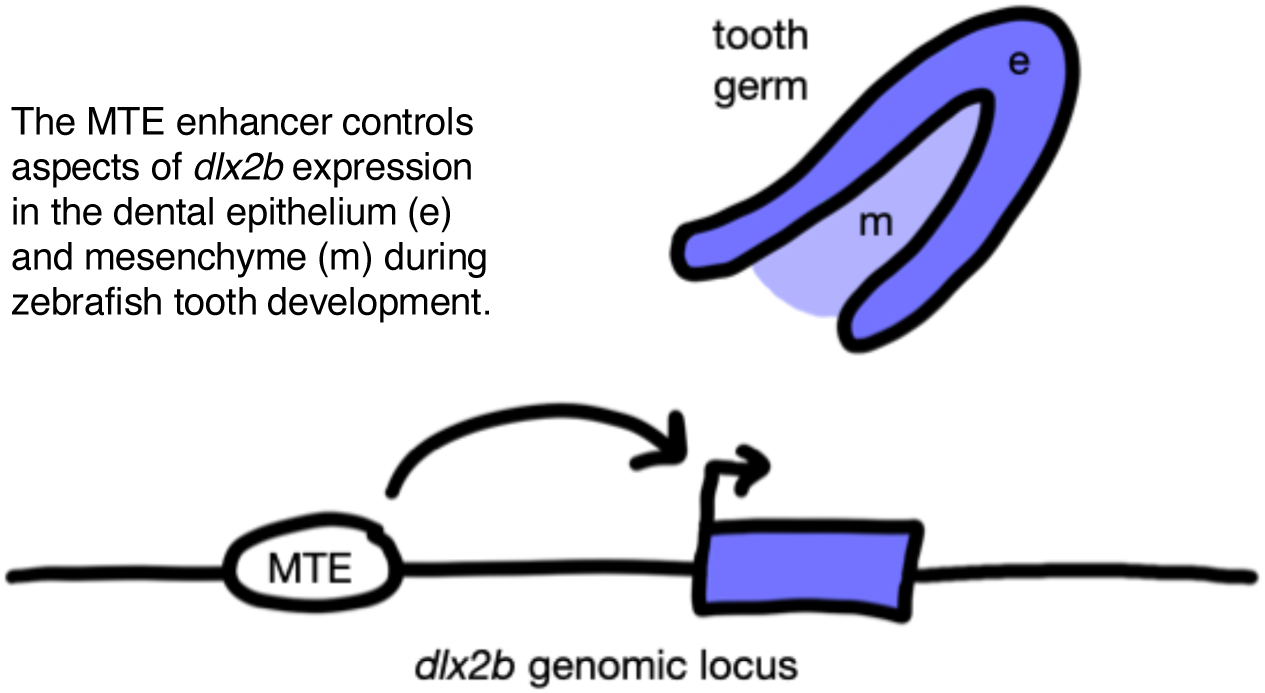

## Introduction

Techniques such as mRNA *in situ* hybridization and reporter transgenics have allowed the visualization of the often-complex spatiotemporal expression patterns of many genes during embryogenesis (e.g. Jensen, 2014; Kvon, 2015). However, the mechanisms behind how the transcription of these genes are regulated to produce complex developmental patterns has been much more difficult to ascertain (Pachano et al., 2022). Now-classic “promoter-bashing” functional identification and analysis of *cis-*regulatory promoters and enhancers provided a foundation for the field (e.g. Goto et al., 1989), and large-scale bioinformatics work continues to be done at the genomic level to identify *cis-*regulatory elements of developmental genes and to determine their function (e.g. Davis et al., 2018). However, there remains a need for functional examination of individual elements to provide data upon which to test genome-wide predictions and to better understand the nature of *cis-*regulation in general, including enhancer/promoter interactions (Oudelaar et al., 2019) and predictions of sequences with specific functions (Grossman et al., 2017). Thus, new, direct tests of specific parts of developmental *cis-*regulatory elements are beneficial for the field as a whole.

Developing tooth germs are a good system in which to examine *cis-*regulatory function, as the expression of a large number of genes has been characterized in these primordia and the anatomy is relatively simple (Balic and Thesleff, 2015; Catón and Tucker, 2009). Additionally, gene expression and function during tooth development has been investigated in a wide range of vertebrate species ranging from mammals to sharks (Jernvall and Thesleff, 2012; Rasch et al., 2016), and these evolutionary comparisons have shown that, especially in early stages of tooth germ formation, vertebrate teeth develop remarkably similarly (Fraser et al., 2009). A large number of different classes of genes have been examined in tooth formation, including transcription factors like Distal-less (Dlx) genes (Borday-Birraux et al., 2006; Zhao et al., 2000), cell signaling molecules like Sonic hedgehog (Shh; Seppala et al., 2017), and structural genes involved in cell behavior and differentiation such as Cadherins (Verstraeten et al., 2013).

However, despite the relatively abundant information regarding gene expression, very little is yet known about the tooth-related *cis-*regulation of most of these genes.

Only a small number of *cis-*regulatory elements capable of driving specific expression patterns during tooth formation have been previously described. Perhaps the most well characterized are enhancers for mouse *Shh* (Sagai et al., 2009; Seo et al., 2018) and the stickleback bone morphogenetic protein gene *bmp6* (Erickson et al., 2015; Cleves et al., 2018; Stepaniak et al., 2021), both of which have been isolated in reporter constructs as well as functionally examined with mutagenesis experiments that have identified potential transcription factor binding sites. Genomic regions near other tooth-related genes have been shown to drive tooth-specific reporter expression, such as from medaka *sp7* (Renn and Winkler, 2009) and human *RUNX2* (in zebrafish; Kague et al., 2012), but the exact position and makeup of enhancers in these regions has not been determined. Similarly, reporters capable of driving tooth-specific expression have been isolated from 4 kb of genomic sequences immediately 5’ of mouse *Dlx2* (Thomas et al., 2000) and zebrafish *dlx2b* (Jackman and Stock, 2006), but again, specific enhancers in these regions have not been identified. Thus, even in the few tooth-related enhancers identified, much remains to be learned about their makeup and function.

Dlx genes are homeodomain transcription factors involved with many aspects of development in vertebrates, especially craniofacial formation and the development of teeth (Jeong et al., 2008). As vertebrate tooth germs begin to form, the most significant tissues involved are the inner dental epithelium, which will eventually differentiate into ameloblasts and secrete the hypermineralized outer part of the tooth (enamel or enameloid, depending on the species), and the dental mesenchyme or papilla, a cranial neural crest derived tissue that will differentiate into odontoblasts and make the bone-like dentin inner layer (Huysseune et al., 1998; Peters and Balling, 1999). Dlx genes are expressed in overlapping spatiotemporal domains of these two developing tissues, with six orthologs activated during both mammalian and zebrafish tooth development (Borday-Birraux et al., 2006; Zhao et al., 2000). Zebrafish *dlx2b* is a particularly approachable gene in the context of tooth development, as it is expressed in a more tooth-specific way relative to neighboring tissues than other Dlx paralogs (Borday-Birraux et al., 2006). Zebrafish develop larval teeth in tissue lining the ventral, posterior part of the pharynx, just anterior to the esophagus (Huysseune et al., 1998). *dlx2b* is expressed in the inner dental epithelium during early morphogenesis stages of tooth development, as well as later in both the epithelium and mesenchyme tissues in differentiating ameloblasts and odontoblasts, respectively, and this transcriptional activity continues until tooth attachment (Borday-Birraux et al., 2006; Jackman et al., 2004). The *cis-*regulation of *dlx2b* has been studied in regard to enhancers located 3’ to the gene which control aspects of its expression during brain development (MacDonald et al., 2010) and, as mentioned above, 4 kb of genomic sequence 5’ to the gene are capable of driving reporter expression during tooth formation (Jackman and Stock, 2006), but little else is known about the function of this 5’ region.

In this study, we have pared down this *dlx2b* 5’ genomic region to a minimal functional enhancer of tooth germ cell reporter expression and performed mutagenesis tests on predicted transcription factor binding sites of interest. Additionally, we have created a *dlx2b* GFP knock-in line that recapitulates the tooth germ expression of the gene and tested the function of the identified minimal tooth enhancer by deleting its core sequences and observing the resulting change in expression. Together, these experiments provide new empirical data regarding the *cis-*regulatory control of tooth organogenesis.

## Results

### Identification of a dlx2b minimal tooth enhancer

Approximately two days prior to the appearance of a given mineralized zebrafish pharyngeal tooth (Fig. 1A), the cells of its tooth germ are already well-organized and transcribing a number of tooth-related genes, including *dlx2b* (Fig. 1B). Tooth germ *dlx2b* expression has been observed in at least the first four zebrafish tooth germs that form (4V^1^, 3V^1^, 5V^1^, and 4V^2^; Borday-Birraux et al., 2006) and thus appears to be a robust marker of odontogenesis.

**Fig. 1.**
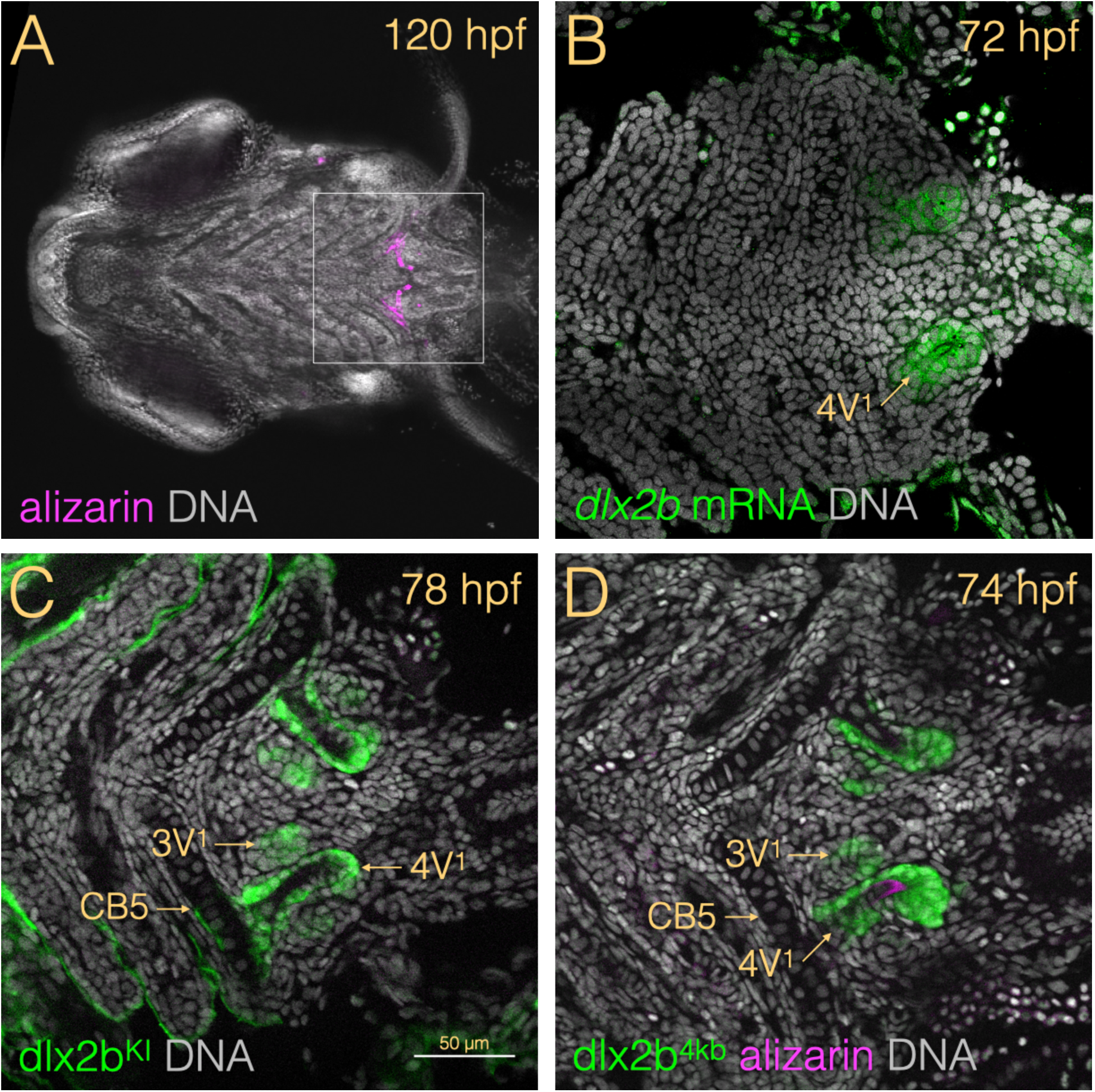
Endogenous and reporter expression of *dlx2b* in developing zebrafish tooth germs. (A) Ventral view with anterior to the left of a 120 hpf larval head labeled with alizarin red S (magenta) to show mineralizing tissues and DAPI (white) to stain nuclei and thus provide a cellular context. The tooth-forming region in the posterior pharynx is indicated (box), highlighting the approximate zoom and cropping of the subsequent panels, in which the right-side (bottom) tooth germs are labeled. (B) Optical section of *dlx2b* mRNA *in situ* hybridization (green) at 72 hpf with the 4V^1^ tooth germ indicated. (C) Expression from the dlx2b^KI^ reporter line showing the 3V^1^ tooth germ at a morphogenesis stage and 4V^1^ at cytodifferentiation. (D) GFP expression in the dlx2b^4kb^ reporter, additionally stained with alizarin red S to show developing tooth mineralization. CB5 = fifth ceratobranchial cartilage (onto which all of the teeth eventually attach).

In order to better understand the *cis-*regulatory control of *dlx2b* during this process we undertook two approaches. One methodology was to create a reporter line driven by the endogenous *cis-*regulatory sequences at the *dlx2b* locus so we could have a way of assessing the full expression pattern of *dlx2b* more easily and at later stages than is feasible using mRNA *in situ* hybridization. Our approach was to make a knock-in (KI) allele at the locus, termed dlx2b^KI^, where the coding sequence for the green fluorescent protein (GFP) along with an attached promoter was inserted using CRISPR/Cas9 (see Methods, Fig. S1; Ota et al., 2016). In individuals heterozygous for this dlx2b^KI^ allele, tooth-related GFP expression is observed in a pattern that appears to recapitulate those seen both by mRNA *in situ* hybridization and previous reporter analysis (Fig. 1C; Jackman & Stock 2006) and tooth germs and subsequent tooth development appears morphologically normal. We hypothesize that dlx2b^KI^ is reporting the normal expression pattern of *dlx2b* in developing tooth germs because the overall expression pattern of *dlx2b* appears to be recapitulated (including early CNS expression (MacDonald et al., 2010); not shown), and this method has reproduced accurate expression patterns for other developmental genes (Kimura et al., 2015; Ota et al., 2016). Similarly to *dlx2b* mRNA (Borday-Birraux et al., 2006), GFP expression from the dlx2b^KI^ allele is observed in the inner dental epithelium in early (morphogenesis) stages of tooth germ formation and at later (cytodifferentiation) stages is present in the dental mesenchyme as well. However, differently than reported for *dlx2b* mRNA, we also sometimes observed a small amount of dental mesenchyme expression in morphogenesis stage tooth germs. This pattern appears to be the same for at least the first four teeth to form (4V^1^, 3V^1^, 5V^1^, and 4V^2^). In the figures presented here, we have chosen to show 3 day post fertilization (dpf, 72-78 hpf) images as representatives, as these often display 3V^1^ in a relatively early/morphogenesis stage in the same focal plane as 4V^1^, which is in a later/cytodifferentiation stage, and thus captures most all of the variation in expression that we observed in tooth germs at various stages. We noticed no tooth-specific differences in expression (e.g. 4V^1^ vs. 3V^1^, 5V^1^, or 4V^2^) for any of the results reported in this study.

The second approach to understand *dlx2b cis-*regulation involved isolating a minimal enhancer from the *dlx2b* locus capable of driving a normal tooth germ expression pattern. As mentioned above, it was previously determined that the 4kb of genomic sequence immediately 5’ of the *dlx2b* transcription start site, when used as a promoter for GFP reporter expression in a transgene allele here referred to as dlx2b^4kb^, is sufficient to drive expression in a *dlx2b*-like pattern (Fig. 1D; Jackman and Stock 2006). Using a promoter-bashing type approach, we tested a series of reporter constructs, first trimming sequences from the 5’ end of the 4kb region, and then, using non-*dlx2b* minimal promoters, from the 3’ end (Table S1). The result was the isolation of a 215 bp region located 817-604 bp 5’ of the *dlx2b* translation start site which was sufficient to drive GFP expression in developing tooth germs, a sequence which we designate as the *dlx2b* minimal tooth enhancer (MTE). This region contains sequence homology with that of other vertebrate genomes, including humans (Fig. 2A), and four regions of very high conservation that may represent conserved transcription factor binding sites (Fig. 2B). These four regions of high homology had been noted previously (Jackman and Stock 2006) and, in retrospect, would have been a good phylogenetic-footprinting type guide to have followed in isolating this enhancer.

**Fig. 2.**
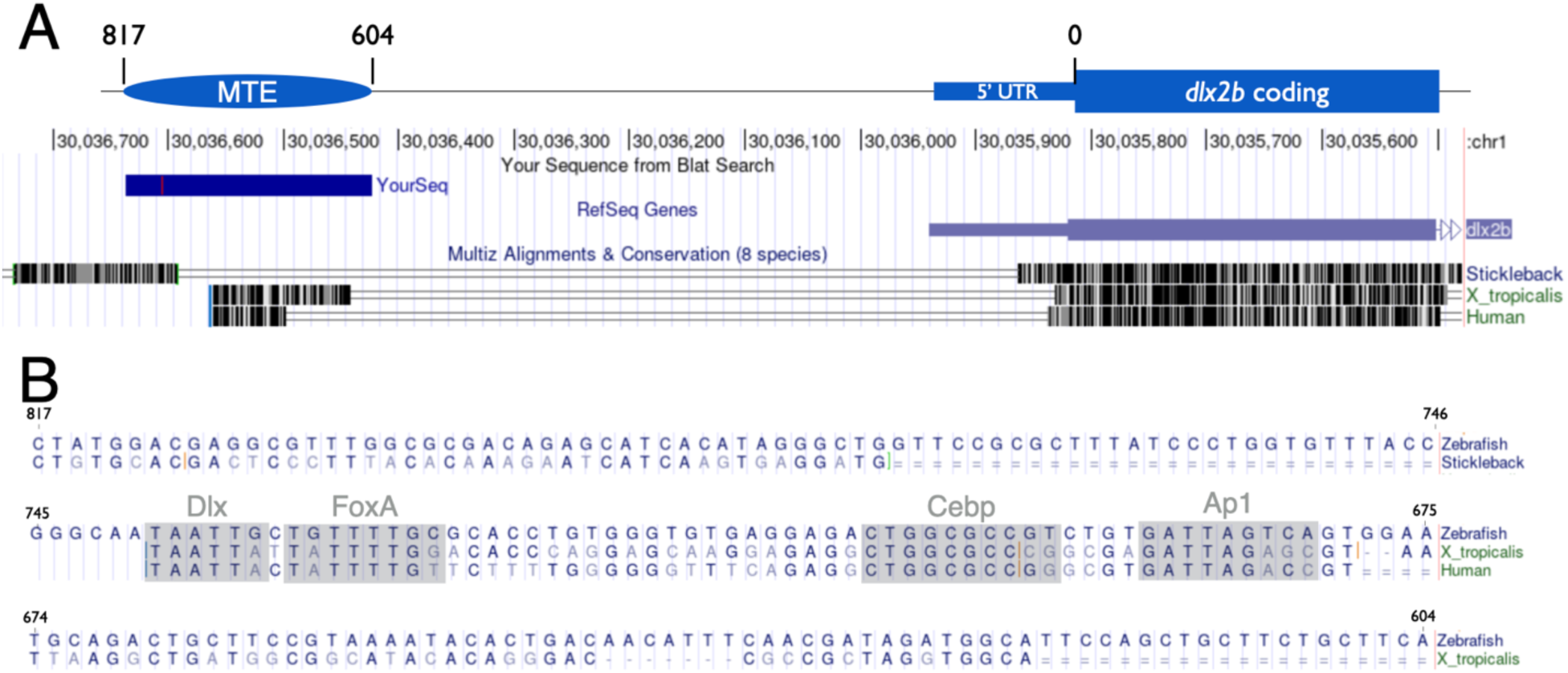
Location and sequence of the *dlx2b* minimal tooth enhancer MTE. **(A)** Diagram of the 5’ end of the zebrafish *dlx2b* gene, including the 5’ UTR and 817 bp of upstream non-coding sequence. The location of the identified 215 bp minimal tooth enhancer MTE is indicated. The black bars below the gene schematic represent regions of homology with the stickleback, *X. tropicalis,* and human genomes (UCSC Genome Browser Zv9/danRer7). (B) Sequence of MTE aligned with those of other species. Predicted possible transcription factor binding sites with high conservation are shaded.

The GFP reporter transgene, termed dlx2b^MTE^, was then established as a stable line. When its expression is compared with the dlx2b^KI^ (Fig. 3A) and dlx2b^4kb^ (Fig. 3B) lines, the GFP expression pattern of dlx2b^MTE^ appears extremely similar (Fig. 3C), with strong early expression in the inner dental epithelium and later expression in the dental mesenchyme. When a smaller region of MTE spanning only the 744-674 bp upstream of the *dlx2b* translation start site was tested in transient assays, no reporter expression was seen (n=13; Fig. 3D), suggesting that there are sequences, possibly other TF binding sites, important for the function of the enhancer on the edges of the 215 bp MTE, and not just within the highly conserved central region that encompasses the four conserved, predicted TF binding site sequences.

**Fig. 3.**
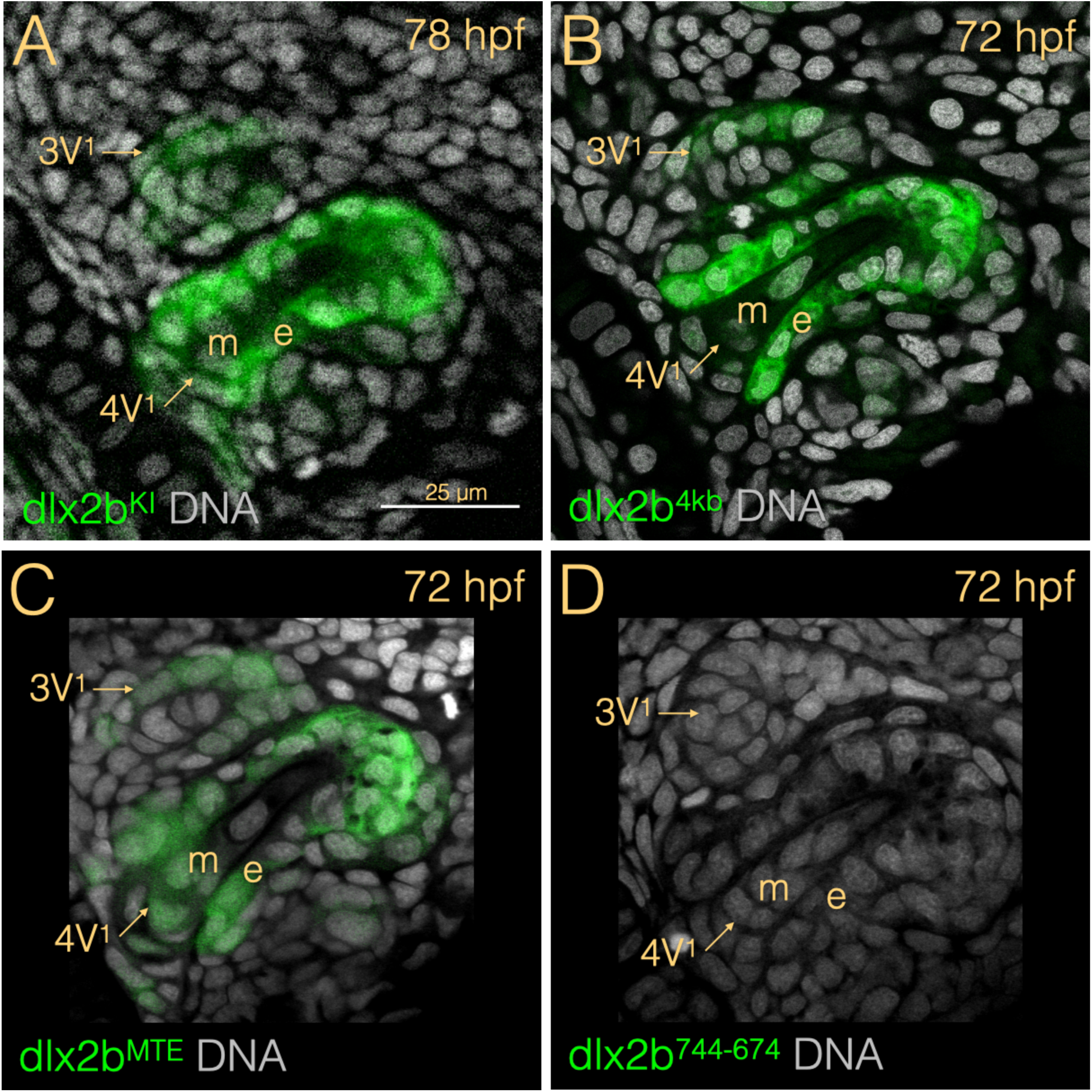
Tooth germ reporter expression. Ventral view close-up of one side of the tooth forming region with the tooth germs 4V^1^ and 3V^1^ indicated. The dlx2b^KI^ reporter line (A), the dlx2b^4kb^ line (B), and dlx2b^MTE^ (C), all exhibit GFP expression primarily in the inner dental epithelium (e) but also somewhat in the dental mesenchyme (m), whereas no GFP expression was observed in the dlx2b^744-674^ truncated reporter (D).

### Characterization of the function of MTE

We next wanted to test if several predicted transcription factor (pTF) binding sites within the MTE region were necessary for tooth germ enhancer activity. Our approach was to create a series of reporter constructs with the pTF binding sites altered so as to maximally disrupt their consensus sequences and test these mutated versions in transient injection assays (Fig. S2; Table S1). We found that none of the six types of pTF sites tested eliminated tooth germ expression when mutated individually (Dlx, FoxA, Cebp, Ap1) or as groups of the same TF type (Pea3/Etv4 or Hox/Pbx/Meis; Fig. S2). However, when all four of the most highly conserved pTF sites were mutated simultaneously (Dlx, FoxA, Cebp, and Ap1; designated DFCA), tooth germ reporter expression was eliminated (n=30; Fig. 4), suggesting that these sites are together necessary for MTE function.

**Fig. 4.**
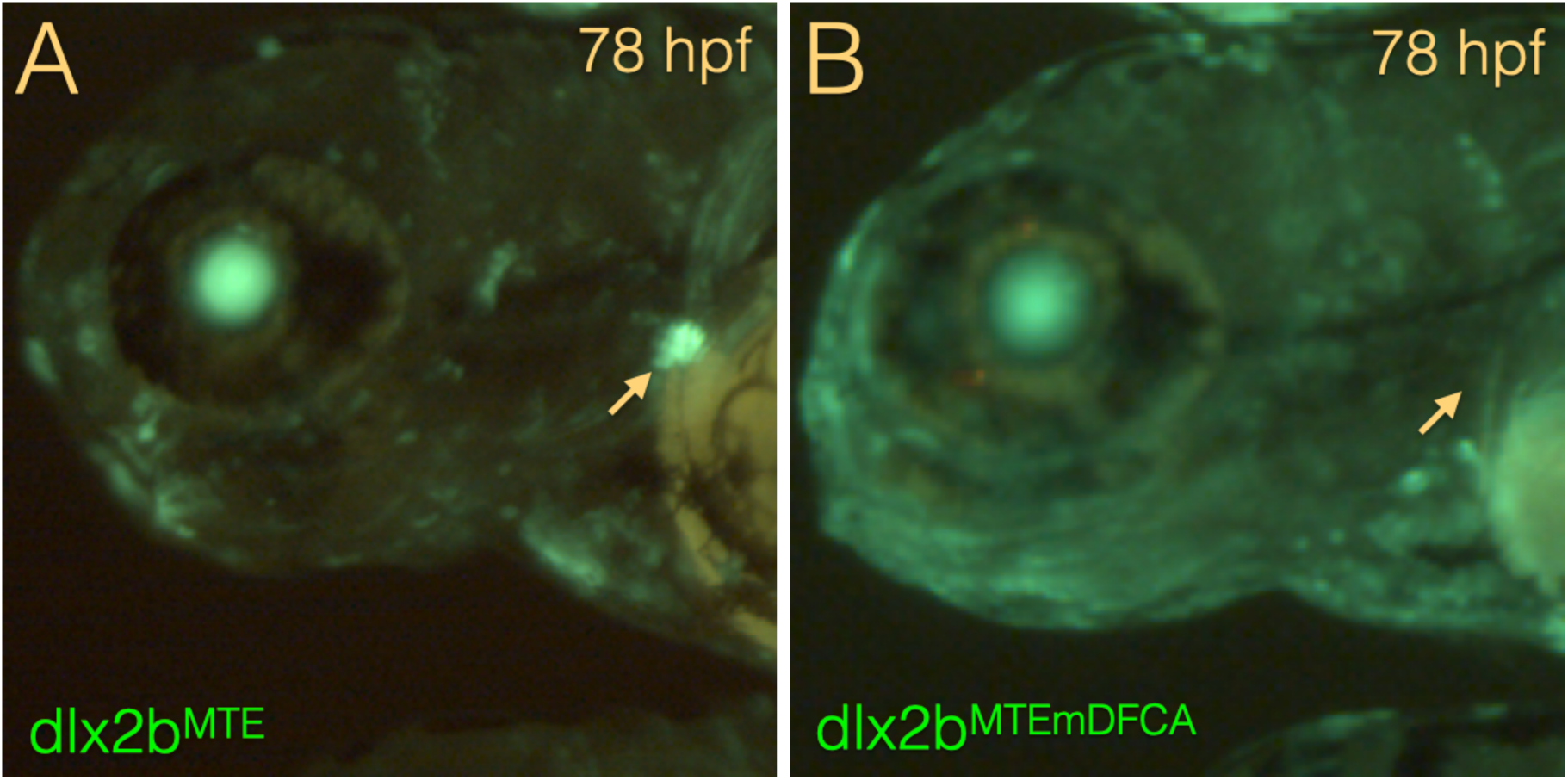
Conserved transcription factor binding sites are required for MTE function. Zebrafish larvae injected transiently with either the dlx2b^MTE^ GFP reporter construct (A) or the dlx2b^MTEmDFCA^ construct with the predicted Dlx, FoxA, Cebp, and Ap1 binding sites all mutated (B). Arrows indicate the location of the tooth-forming region.

Because of the apparent importance of the conserved DFCA pTF sequences in the MTE, we examined the tooth-related mRNA expression of some of the likely genes whose proteins may be binding to these sequences during morphogenesis and cellular differentiation stages of 4V^1^ tooth germ formation (Fig. S3). We observed the expected 4V^1^ tooth germ expression of two Dlx genes (Jackman et al., 2004; Borday-Birraux 2006), but found no apparent tooth-germ-related expression for *foxa2*, which had previously been reported expressed near the tooth-forming pharyngeal region (Piotrowski and Nüsslein-Volhard, 2000), or the *jun* Ap-1 subunit gene (Fig. S3). In contrast, we did observe tooth related expression for *cebpa*, which upon closer examination using fluorescence mRNA ISH and a GFP knock-in line constructed similarly to dlx2b^KI^, was seen to be localized mostly to the inner dental epithelium in a domain near the tooth base, overlapping a part of the *dlx2b* expression pattern (Fig. S4). Thus, gene expression analysis reveals that Dlx and Cebp genes may be particularly important for activating *dlx2b* expression in at least the developing 4V^1^ tooth germ.

Next, we wanted to test the necessity of the *dlx2b* MTE region for regulating normal tooth gene expression. To do this in a feasible way, rather than targeting an unaltered *dlx2b* locus, we used CRISPR/Cas9 to create a deletion within the MTE in the dlx2b^KI^ allele so that we would be able to immediately gauge any resulting changes to expression via GFP. Using two guide RNAs, we deleted 88 bp within the *dlx2b* MTE, generating an allele termed dlx2b^KIΔMTE^ (Fig. S4). This deletion spans from 760 to 673 5’ of the *dlx2b* translation start site and eliminates all four of the highly conserved pTF biding sites Dlx, FoxA, Cebp, and Ap1 (Fig. 2B). The mutation had a major effect on the tooth germ GFP expression pattern, seemingly reversing the pattern relative to the dental epithelium and mesenchyme (Fig. 5). Whereas normally dlx2b^KI^ GFP expression is observed during morphogenesis and cytodifferentiation stages mostly in the inner dental epithelium with lower-appearing levels of expression in the dental mesenchyme (Fig. 5A), in the dlx2b^KIΔMTE^ allele, GFP is strongly expressed in the dental mesenchyme (Fig. 5B) with weaker, occasionally observed expression in the distal part of the inner dental epithelium (not shown). Thus, it appears likely that in the *cis-*regulatory context of the genomic locus surrounding *dlx2b*, the MTE has an important function in directing correct tissue-specific expression within developing tooth germs.

**Fig. 5.**
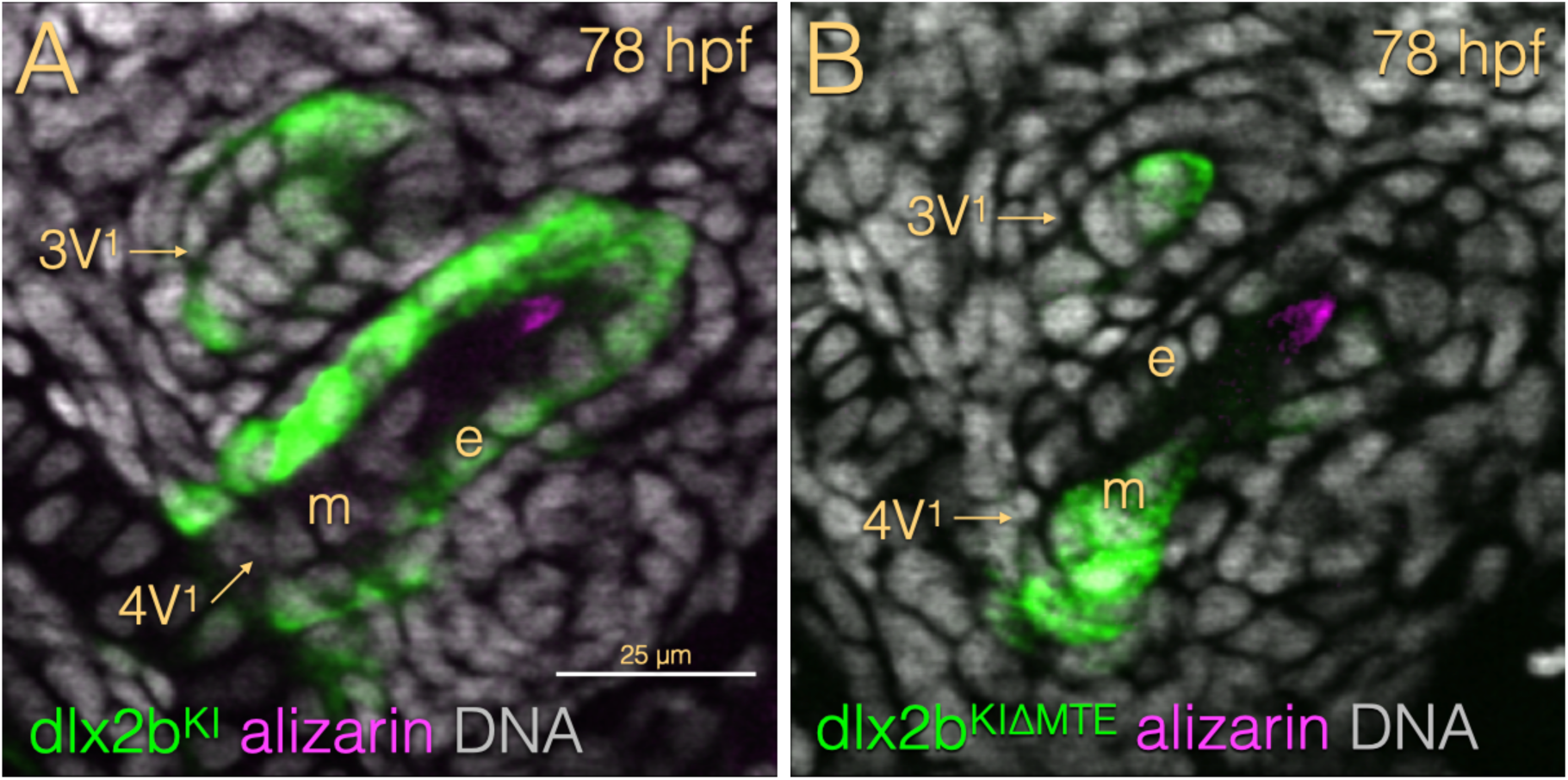
MTE is required for proper tissue-specific expression in developing tooth germs. (A) In dlx2b^KI^ at 78 hpf, GFP expression appears much stronger in the inner dental epithelium (e) than in the dental mesenchyme (m) in both 4V^1^ and 3V^1^ tooth germs. (B) In dlx2b^KIΔMTE^ at the same stage, the expression pattern appears reversed, with strong GFP signal in the dental mesenchyme and little to none in the epithelium.

## Discussion

We have shown that the *dlx2b* minimal enhancer MTE is sufficient to recapitulate most or all of the normal *dlx2b* expression pattern in early developing zebrafish tooth germs, especially that of the inner dental epithelium. MTE contains four regions of conspicuously high sequence conservation that are predicted to represent important transcription factor binding sites (pTFs), but this region by itself, as shown in the dlx2b^744-674^ reporter experiment, is not capable of driving tooth-specific expression. Additionally, no one of these conserved sites seems necessary for the overall activity of the enhancer, suggesting some redundancy in their function, either with each other or with sequences elsewhere in the genome. More work will be required for firm answers regarding what each pTF site might specifically control with regard to *dlx2b* transcriptional activity, but it is nevertheless interesting to consider a few ideas of how they might function. For example, FoxA proteins have been shown to sometimes act as pioneer transcription factors, opening up chromatin regions for further transcription factor binding (Iwafuchi-Doi and Zaret, 2016). Although, our analysis did not detect expression of one zebrafish FoxA paralog, *foxa2*, in the tooth germ of 4V^1^ (Fig. S3), this gene has been shown to be expressed in the earlier pharyngeal tooth-forming region of zebrafish (Piotrowski and Nüsslein-Volhard, 2000), consistent with the idea that the FoxA pTF site in MTE might be important in chromatin modification, perhaps at early stages before the first-forming tooth germ is distinguishable. Another example is that of Cebp proteins, which have been studied specifically with regard to mouse tooth development and are known to be important in ameloblast differentiation and activation of amelogenin transcription (Zhou and Snead, 2000; Xu et al., 2007). We observed *cebpa* expression in a region of the zebrafish tooth germ inner dental epithelium consistent with these roles, and in a pattern which seems to be a subset of the full *dlx2b* epithelial expression domain (Figs. S3 and S4), suggesting that *cebpa* could possibly be an activator of *dlx2b* in these cells acting through the MTE Cebp pTF site. Even though the function of these conserved sequences remains unknown, the activity of MTE was eliminated when all four sites were mutated (Fig. 4), emphasizing that these evolutionarily conserved sequences are important and that they may work together, even in isolation from the *dlx2b* locus.

However, it is important also to consider how the MTE enhancer might be working in its endogenous locus and how this could be different than how it works in isolation. In this context, we have observed that in both the randomly inserted transgenes dlx2b^4kb^ and dlx2b^MTE^, GFP reporter expression appears more restricted to the dental epithelium, especially in early morphogenesis stages (Figs. 3B, 3C), when compared with the GFP knock-in allele dlx2b^KI^ (Figs. 3A, 5A). If this observation is accurate, and dlx2b^KI^ indeed does more precisely represent the real expression pattern of *dlx2b* (mRNA *in situ* hybridization isn’t much help in determining what is real, as we find it to be less sensitive than GFP reporters, it is difficult to do at later stages, and the visualization products tend to diffuse), it suggests that there is a small amount of dental mesenchyme transcriptional activity that is not controlled by MTE, and is instead activated by something else. One explanation for this could be the presence of one or more enhancers located at a different location at the *dlx2b* locus that can drive dental mesenchyme transcriptional activation. This idea is consistent with the results from the dlx2b^KIΔMTE^ allele where the evolutionarily conserved core of MTE is deleted and most expression is lost in the dental epithelium, but dental mesenchyme expression appears increased (Fig. 5B). While it is possible that this deletion is somehow changing the MTE from being a mostly-epithelial enhancer into a mostly-mesenchymal one, given the likelihood of other enhancers being in the vicinity, we prefer the following hypothesis. In a normal situation, MTE is interacting with other enhancers and the *dlx2b* promoter, and perhaps due to its proximity to the promoter largely wins, resulting in an expression pattern in the dental epithelium that mirrors what the MTE reporter generates in genomic isolation. However, when MTE is compromised, other enhancers at the *dlx2b* locus more easily interact with the promoter and their expression patterns are increased (Fig. 6). This hypothesis is inspired by a number of studies on enhancer/promoter interactions (e.g. Bateman et al., 2021; Oudelaar et al., 2019). Additionally, while not identical, it is interesting to compare the regulation of *dlx2b* with one of the few other zebrafish tooth developmental genes where *cis*-regulation has been carefully examined: *bmp6*, which also exhibits complex patterns of expression in the developing dental epithelium and mesenchyme (Cleves et al., 2018, Stepaniak et al., 2021).

**Fig. 6.**
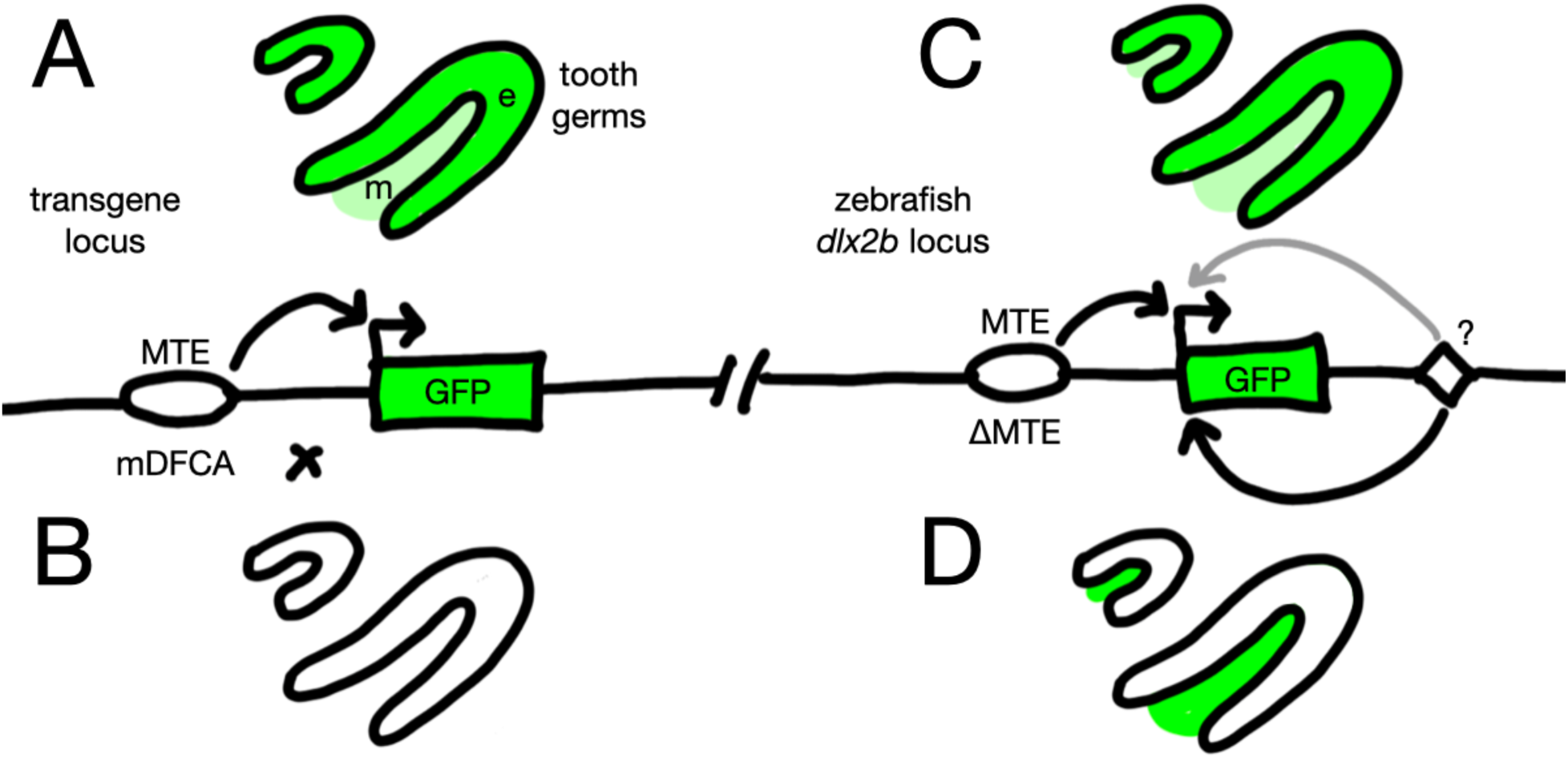
Summary of results and model of possible MTE enhancer function. The left side (A, B) represents the Tol2-based transgene experiments at a remote locus and the right side (C, D) the knock-in experiments at the *dlx2b* locus. The top of the diagram (A, C) depicts the normal MTE sequence and the bottom (B, D) the mutated versions. MTE is sufficient to drive a mostly-epithelial tooth germ expression pattern even at a remote locus (A) but when mutated at a remote locus (B), there are perhaps no other nearby enhancers to compensate and thus all tooth germ expression is lost. In contrast, at the *dlx2b* locus, MTE may be a primary driver of epithelial tooth germ expression (C), but when mutated, one or more other *cis-*regulatory elements (diamond) maintain expression, but in a more mesenchymal pattern (D).

As a caveat, our model relies on an assumption that the dlx2b^KI^ allele with its introduced promoter is behaving like a normal *dlx2b* gene, and thus more work will need to be done both to specifically identify other enhancers at the *dlx2b* locus and to understand how the *cis-*regulatory elements may be together interacting with the endogenous *dlx2b* promoter to generate its complete expression pattern during tooth formation. Some of this information may be acquired along with genome-wide modeling of *cis-*regulatory function based on chromatin structure and other features (e.g. ENCODE; Boyle et al., 2014) but there is also still much that can be learned from targeted functional studies of *cis-*regulation such as this one.

## Methods

### Animal husbandry and anatomy

All experimentation with the zebrafish (*Danio rerio*, Hamilton 1822) in this study followed protocols approved by the Institutional Animal Care and Use Committee at Bowdoin College and at the University of Mississippi Medical Center. The zebrafish used were from an in-house strain, derived originally from a mixture of the Tü and AB lines. Embryos were raised in 30% Danieu’s embryo medium, with the addition of 0.002% methylene blue to reduce fungal growth and kept at approximately 28.5°C. Embryonic stages are reported in hours or days post-fertilization or using the nomenclature of Kimmel et al. (1995).

Individual teeth are labeled following the convention of Van der heyden & Huysseune 2000, with the first-forming tooth designated 4V^1^, and the subsequently forming teeth as 3V^1^, 5V^1^, and 4V^2^ (the replacement of 4V^1^). These teeth develop in a reliable temporal progression as well as in characteristic locations and were thus possible to unambiguously identify. Names for tooth germ developmental stages are as in Huysseune et al. (1998): initiation (not shown in this study), morphogenesis, cytodifferentiation, and attachment (also not shown).

### DNA constructs and transgenic lines

**Table 1.** Transgenic lines used in this study.

| Name | Official Designation | Insertion | Enhancer(s) | Promoter | Reporter |
| --- | --- | --- | --- | --- | --- |
| dlx2b<sup>4kb</sup> | cs1Tg with construct Tg(dlx2b:EGFP) | unknown via Tol2 | dlx2b 5' 4 kb | dlx2b endogenous | GFP |
| dlx2b<sup>MTE</sup> | bo1Tg with construct Tg(dlx2b-actb2:EGFP,myl7:GFP) | unknown via Tol2 | dlx2b MTE | β-actin | GFP |
| dlx2b<sup>KI</sup> | bo2Tg with construct Tg3(hsp70l:EGFP) | dlx2b locus | dlx2b endogenous | hsp70 | GFP |
| dlx2b<sup>KIΔMTE</sup> | bo3Tg with construct Tg3(hsp70l:EGFP) | dlx2b locus | dlx2b endogenous | hsp70 | GFP |
| cebpa<sup>KI</sup> | bo4Tg with construct Tg3(hsp70l:EGFP) | cebpa locus | cebpa 5' endogenous | hsp70 | GFP |

The GFP knock-in alleles dlx2b^KI^ and cebpa^KI^ were generated via CRISPR/Cas9 using a combination of the methods described in Ota et al., 2016 and Burger et al., 2016. The logic of this method is to insert a plasmid at the target locus containing the Hsp70 promoter situated properly adjacent to the coding sequence of GFP so that a precise integration event is not necessary but nearby enhancers will still be able to interact with the introduced promoter (Ota et al., 2016). Briefly, zebrafish embryos were injected into the blastomeres at early 1-cell or 2-cell stages with about 1 nl of a ribonucleoprotein (RNP) mixture of high-fidelity Cas9 protein (Integrated DNA Technologies, IDT), a guide RNA targeting the desired genomic locus (GCCAAGGCTATCCAGAACAG for *dlx2b*, custom synthesized by IDT), a plasmid containing the Hsp70 promoter and GFP coding sequence (Mbait-hs-eGFP; Ota et al., 2016), a second guide RNA to linearize the plasmid, 300 mM KCl to promote RNA solubility (Burger et al., 2016), and a trace amount of phenol red to facilitate visibility of the solution during injection. As they developed, injected (F0) embryos were scored for GFP expression patterns that matched previously reported mRNA expression, and individuals with correct-looking patterns and relatively low mosaicism were raised as potential founders. Approximately 5% of injected F0 fish showed promising GFP expression and of those, about 25% had germline transmission. The dlx2b^KI^ allele is officially designated in the Zebrafish Information Network (ZFIN) as genomic feature bo2Tg with construct Tg3(hsp70l:EGFP). The complete insert region was PCR amplified using Q5 Hi Fidelity DNA polymerase (NEB) and the amplicon sequenced at high coverage using Illumina MiSeq in the Complete Amplicon Next-Generation Sequencing service from the MGH CCIB DNA Core (Fig. S1).

The dlx2b^KIΔMTE^ allele was made by injecting dlx2b^KI^ embryos with two additional sgRNAs targeting either side of the MTE highly conserved region (CTGGTTCCGCGCTTTATCCC and CGCCGTCTGTGATTAGTCAG) and scoring F0 fish for reduced levels of GFP expression (Fig. S4). Once a stable line was isolated, the deletion region was PCR amplified with Q5 polymerase and characterized via Sanger sequencing. This dlx2b^KIΔMTE^ allele is designated on ZFIN as genomic feature bo3Tg with construct Tg3(hsp70l:EGFP).

The cebpa^KI^ allele is designated on ZFIN as genomic feature bo4Tg with construct Tg3(hsp70l:EGFP) and was created using GGCGGGTTTTAGATACTCCA as a guide RNA. This allele has not been sequenced, but the plasmid insertion into the *cebpa* locus has been verified by diagnostic PCR. The plasmid is in the reverse orientation (hsp70 promoter and GFP towards the 3’ end of the gene) and there is deletion of unknown size 3’ of the plasmid insertion.

The allele we refer to in this study as dlx2b^4kb^ is designated as genomic feature cs1Tg with construct Tg(dlx2b:EGFP). It consists of 4 kb of genomic sequence, 5’ to the translation start site of the *dlx2b* gene, including the endogenous promoter for *dlx2b*. DNA reporter constructs representing subsets of this original 4 kb region were created during three different time periods of the project, each period using somewhat different methods. The parts of each construct are summarized in Table S1. In the first period, constructs were assembled using the Gateway Tol2kit (Kwan et al., 2007), employing the pDestTol2CG2 destination vector, which flanked the insert regions with Tol2 transposon inverted repeats for efficient genomic integration (Kawakami, 2007). The insert region of these plasmids consisted of a fragment of the *dlx2b* 5’ genomic *cis-*regulatory region that sometimes included the *dlx2b* endogenous promoter, 500 bp of the ß-actin promoter (Higashijima et al., 1997) if the *dlx2b* promoter was not included, and EGFP coding sequence with a SV40 polyadenylation signal. In the second period of reporter construction, identically arranged insert regions were created (more quickly and reliably) using NEBuilder (New England Biolabs) and PCR amplified using the Q5 proofreading polymerase (New England Biolabs) without cloning into a plasmid. Directly testing PCR products in this manner was rapid but injection of linear DNA produced a high frequency of deformed embryos that reduced the efficiency of the method. Thus, in the most recent period, a final plasmid construct was made using NEBuilder with a pUC19 plasmid backbone, this time using the hsp70 promoter instead of ß-actin, which decreased non-specific GFP expression in injected embryos (Erickson et al., 2015). All constructs, including PCR products, were verified by Sanger sequencing.

To test the function of predicted transcription factor binding sites, mutations were designed to maximally disrupt the predicted transcription factor binding, while at the same time altering a small number of actual nucleotides to make it more likely that we were only disrupting one binding site. To achieve this balance, we examined the binding matrixes for each transcription factor (Cartharius et al., 2005) and changed all nucleotide positions that were invariant or highly probably associated with a functional binding site (Fig. S2).

Individual F0 embryos injected with these reporter constructs displayed variable expression in tooth germs, if any expression was seen at all, as is expected from mosaic integration of DNA injected into zebrafish embryos (Ni et al., 2016). Despite this mosaicism, because of the lack of potentially confounding nearby expression driven by the *dlx2b cis-*regulatory regions tested, it was unambiguous when a particular construct was driving tooth-related expression, even if it was present in only a subset of cells in the developing tooth germs. Nevertheless, for certain constructs we wanted to observe in detail the non-mosaic pattern of a stably integrated reporter line, and thus raised these fish to at least the F2 generation (Table S1). Due to housing space limitations, most of these stable lines were discarded after F2 testing, but we have retained the line created from construct #9 (Table S1), which we refer to here as dlx2b^MTE^, and is officially designated in ZFIN as genomic feature bo1Tg with construct Tg(dlx2b-actb2:EGFP,myl7:GFP).

### Histology

For NBT/BCIP mRNA *in situ* hybridization detection, partial mRNA was cloned in the TOPO II dual promoter vector, RNA was synthesized with the appropriate RNA polymerase (SP6 or T7), and the hybridization was performed as previously described (Gibert et al., 2006). Sources for riboprobes: *cebpa* (Fraher et al., 2015), *dlx2a* and *dlx2b* (Gibert et al., 2015), *AP-1/jun* amplified with primers GTTTCTGTGTCCCAAGAACG and ATGTAGCTCAGAAGTCAGGC, *foxa2* with primers ACGAGAAAGAGGCGATCAAT and GATGGTTCACAATGCAAGCT. We first confirmed the detection of the expression of *dlx2b* in the 4V^1^ tooth germ by conventional *in situ* hybridization using NBT/BCIP detection. Based on the literature and on our experience (Gibert et al., 2010) we used 56 hpf as a developmental time point when *dlx2b* mRNA detection is the strongest in the tooth germ (Fig. S3A,B). As previously reported (Borday-Birraux 2006), *dlx2b* mRNA detection by NBT/BCIP decreases strongly after 68 hpf and becomes almost undetectable in 4V^1^ by 72 hpf (Fig. S3C/D), before appearing again in 3V^1^ and 5V^1^ (Borday-Birraux et al 2006). Therefore, we used these two timepoints to assess the expression of TFs predicted to bind to the DFCA sequence of the dlx2b^MTE1^ enhancer.

For fluorescence mRNA *in situ* hybridization, riboprobes were created from custom-synthesized DNA templates matching parts of the genes of interest (gBlocks from IDT). For *cebpa*, the sequence used to make the riboprobe started 685 bp from the translation start site and extended 1000 bp into the 3’ UTR (GenBank:BC056548). This was used as template for PCR, in which a T3 RNA polymerase binding site for transcription of the antisense probe was added as part of the reverse primer. For *dlx2b*, the gene-specific sequence consisted of the first 474 bp of the 3’ UTR (GenBank:BC134899) and the T3 site was synthesized directly into the gBlock template. Synthesis and purification of probes was performed as previously described (Thisse and Thisse, 2008). Hybridization and developing steps were performed as described in Talbot et al. (2010), except that digoxigenin-labeled riboprobe was detected with an anti-digoxigenin antibody conjugated to alkaline phosphatase and visualized with a Fast Red reaction (Lauter et al., 2011), and that DAPI or Sytox Green (depending on the microscope to be used later) was added to the overnight antibody incubation in order to simultaneously observe cell nuclei for orientation purposes.

GFP antibody labeling and alizarin red S staining of mineralized teeth was performed as in Yu et al. (2015). We found that we could combine these two methods in a single specimen (e.g. Fig. 1D; Fig. 5) if the alizarin staining was performed after the antibody label and visualized relatively quickly (within 1-2 days).

### Microscopy & Image Processing

Photographs of head and tooth GFP expression in living zebrafish embryos were taken with a Leica MZ16F stereoscope with a DCF300FX camera. Brightfield imaging of NBT/BCIP *in situ* hybridization was done using a Zeiss Axio Imager with a Zeiss color705 digital camera. For close-up photographs of fixed tissue, to get a clear view of the tooth-forming region, the heart and yolk of embryos or larvae were manually removed with insect pins and the specimens were mounted ventral side up in glycerol under a small coverslip, as previously described (Yu et al., 2015). Z-stacks for 3D image analysis were taken using either a Zeiss 510 Meta laser scanning confocal microscope, a Leica TCS SP8 confocal microscope, or a Zeiss Axio Imager M2 with an Apotome 2 structured illumination attachment. Brightness and contrast levels were adjusted uniformly across each image using FIJI/ImageJ (Schindelin et al., 2012). For most images, a single Z-slice is shown, but for certain specimens (Fig. 1A, Fig. 5) it was desirable to show a thicker Z representation. For these, the stacks were rendered in 3D and visualized with FluoRender (Wan et al., 2017). Colors for fluorescence images were selected to facilitate visibility for diverse vision types (Wong 2011). Final figures were assembled using Keynote (Apple Inc.).

### Bioinformatics

Visualization of evolutionary sequence conservation at the *dlx2b* genomic locus was done using the Jul. 2010 (Zv9/danRer7) zebrafish genome assembly in the UCSC Genome Browser (Kent et al., 2002) and guide RNA targets were chosen with help from the CRISPR/Cas9 Sp. Pyog. target sites track. Predicted transcription factor binding sites were located using a combination of PROMO (Messeguer et al., 2002) and MatInspector (Cartharius et al., 2005). DNA sequence analysis and Figs. S1 and S4 were done with Geneious R11 and Geneious Prime 2022.0.1.

## End Matter

### Author Contributions & Notes

W.R.J. designed research; W.R.J., Y.M., D.R.A., A.A.DF., V.M.N., S.Y.L., E.H.C., H.E.L., E.K.R., A.L.J., C.K.C. and Y.G performed research, collected, and analyzed data; W.R.J. and Y.G wrote the paper. The authors declare no conflict of interest.

## Acknowledgements

We thank Atsuo Kawahara for graciously sending us the GFP knock-in plasmid.

## Supplemental Information

**Fig. S1.**
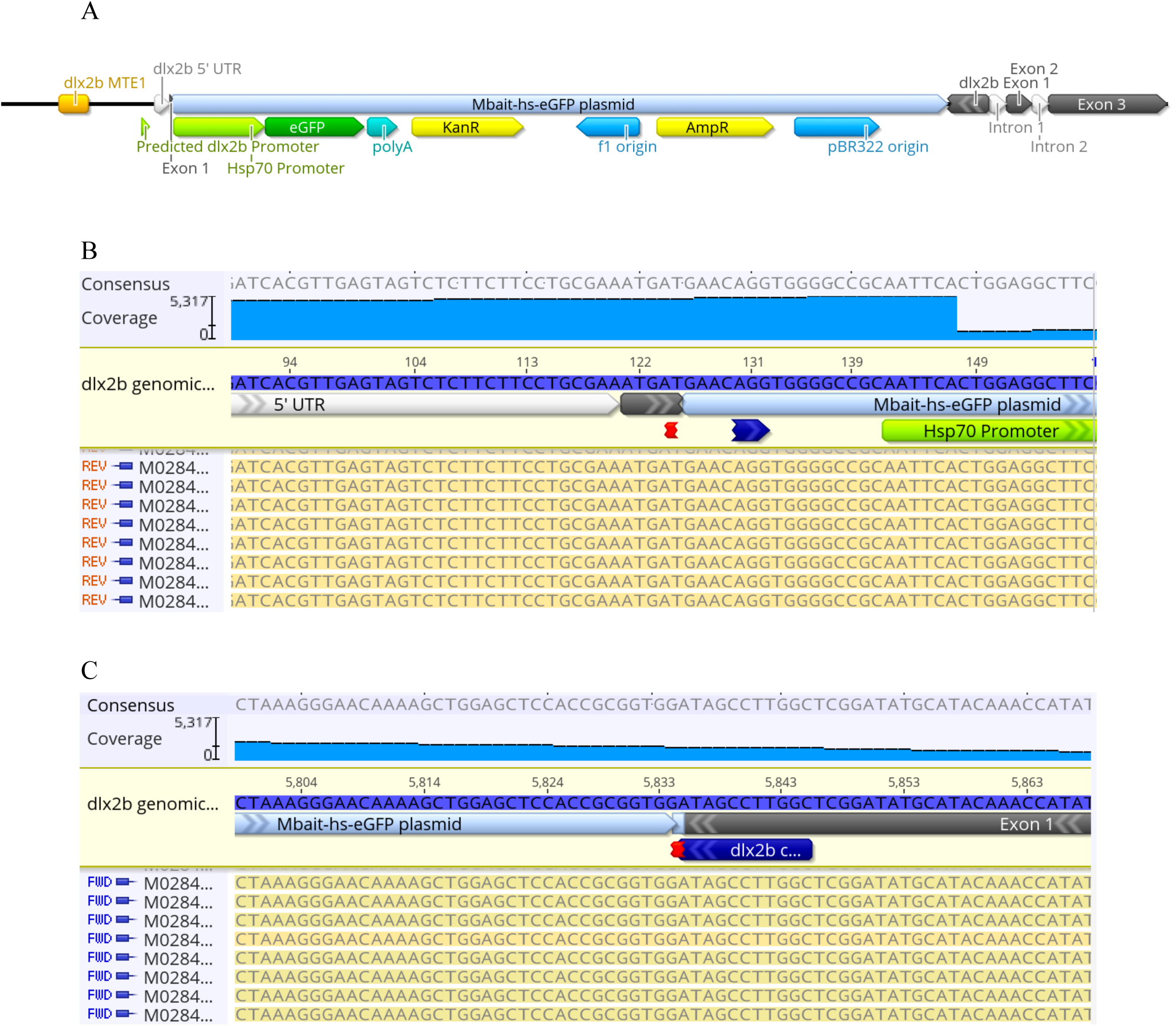
The dlx2b^KI^ allele. (A) Schematic diagram of the Mbait-hs-eGFP plasmid insertion into the *dlx2b* locus. Sequences at the 5’ (B) and 3’ (C) end of the insertion.

**Figure S2.**
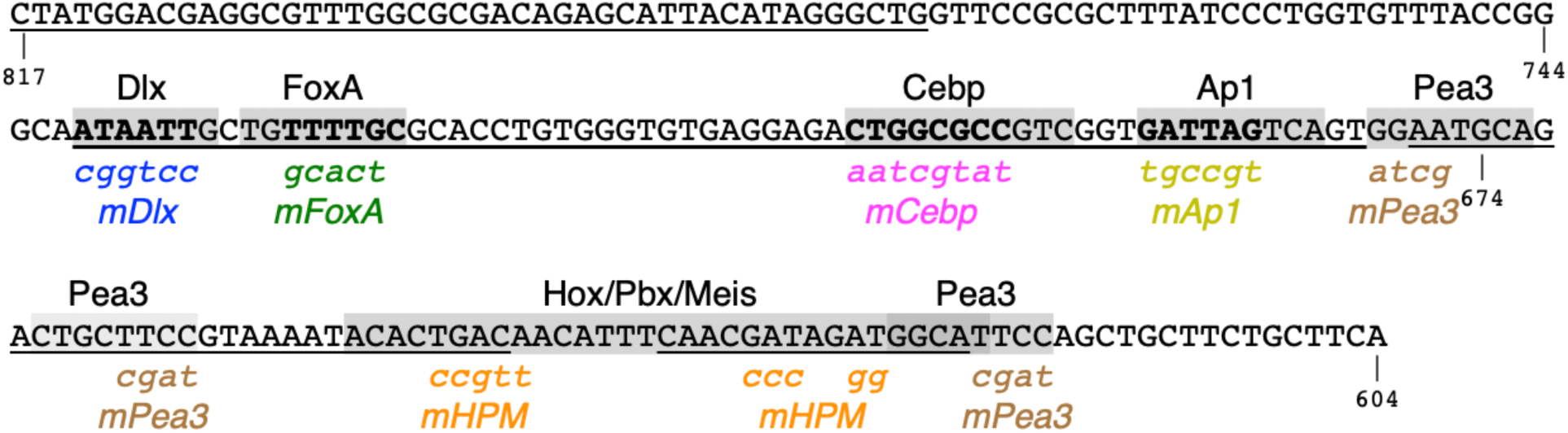
Sequences mutated to test the necessity of various predicted/possible transcription factor binding sites.

**Figure S3.**
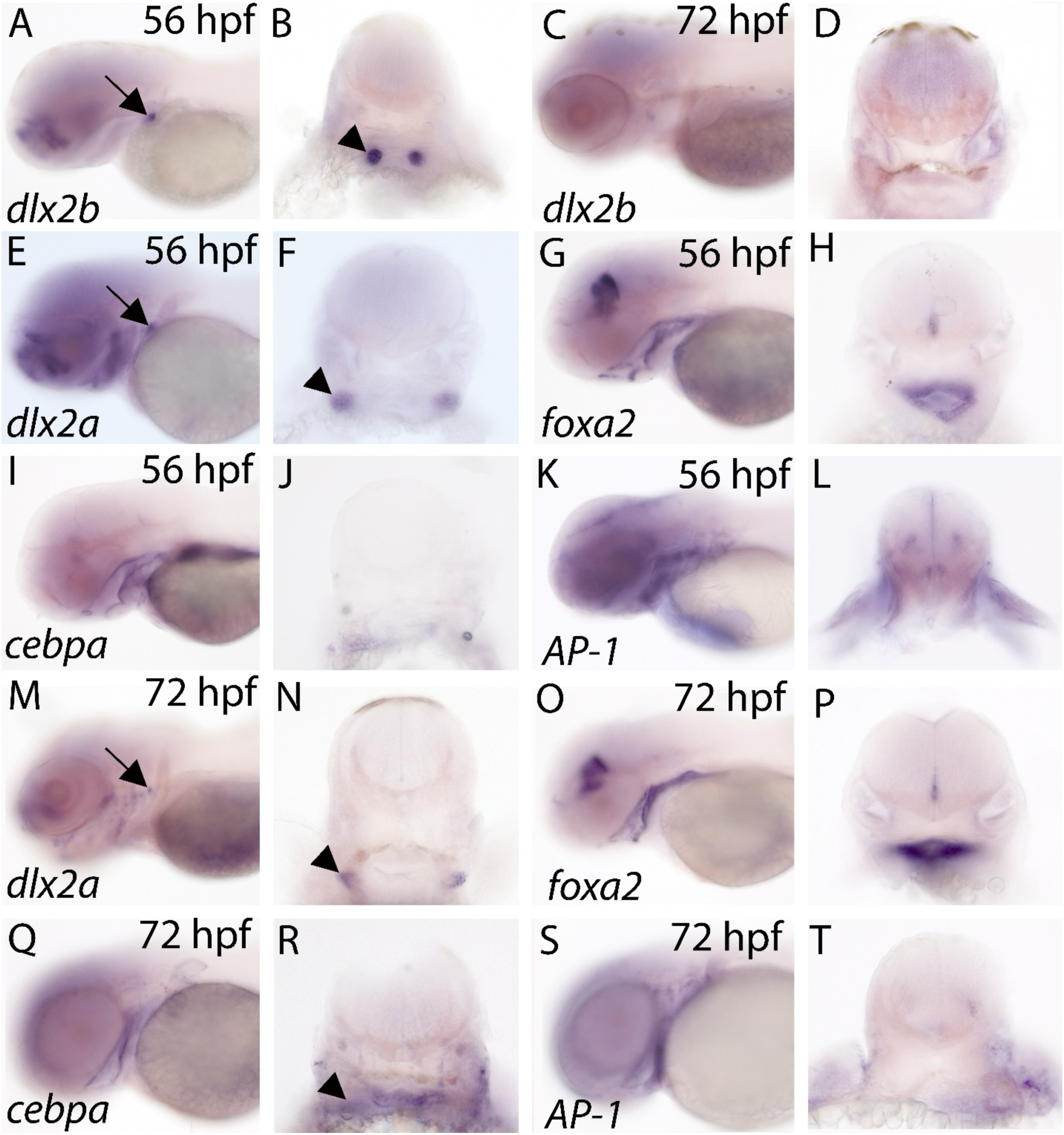
Expression of TFs in and near 4V^1^. (A-D) expression of dlx2b in 4V1 (arrow) at 56 (A) and 72 hpf (C). (B,D) Transverse sections of A and C respectively. The arrowhead in B points to the 4V^1^ tooth germ. (E-L) expression of *dlx2a* (E,F); *foxa2* (G,H); *cebpa* (I,J) and *AP-1/jun* (K,L) at 56 hpf. The arrow in E and the arrowhead in F denote the expression of *dlx2a* in 4V^1^ and more lateral mesenchyme (M-T) expression of *dlx2a* (E,F); *foxa2* (G,H); *cebpa* (I,J) and *AP-1/jun* (K,L) at 72 hpf. The arrow in M and the arrowhead in N denote the expression of *dlx2a* in 4V^1^ and lateral mesenchyme. The arrowhead in R points to the location of expression of *cebpa* in the 4V^1^ tooth germ.

**Figure S4.**
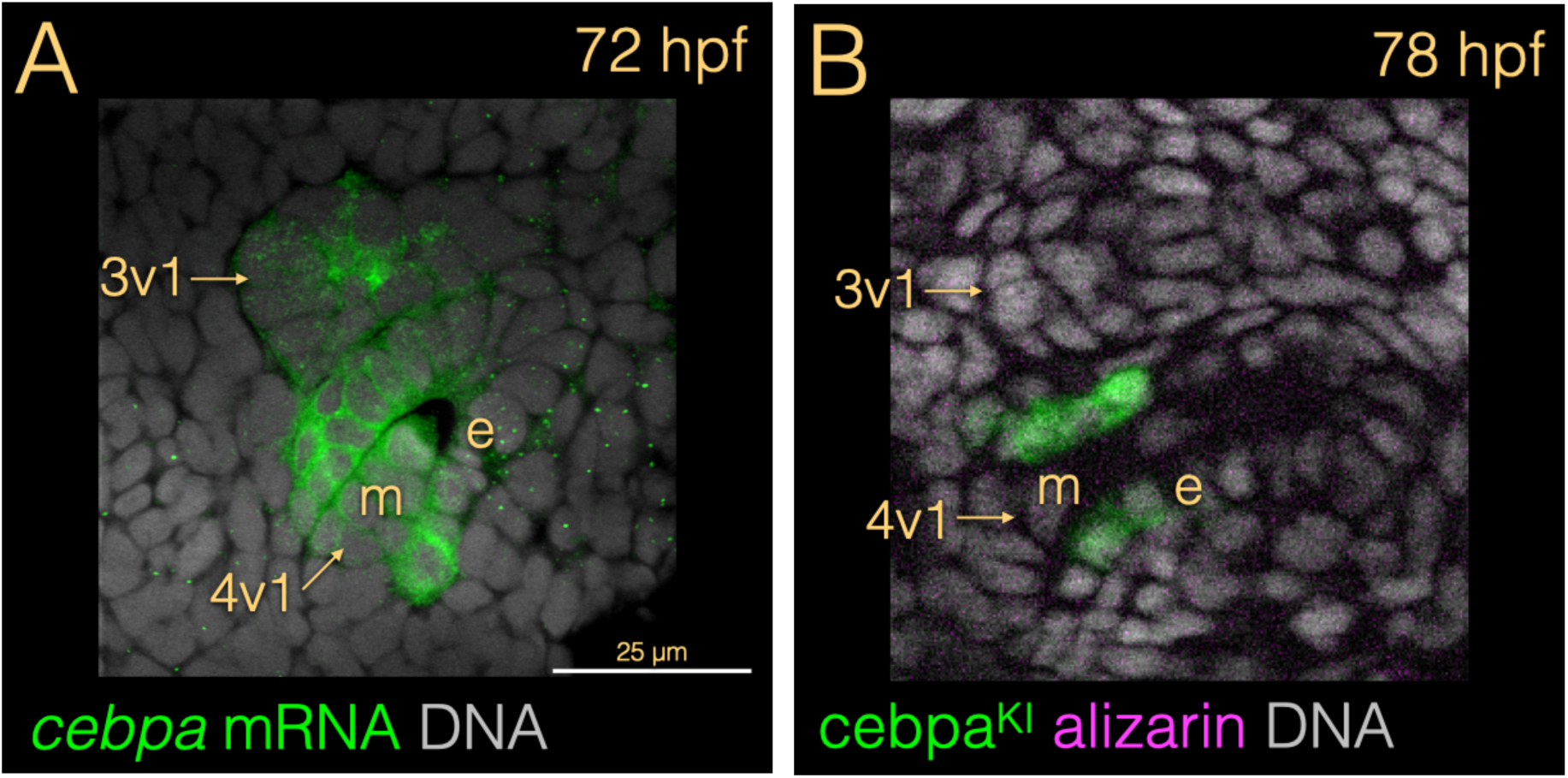
Expression of *cebpa*. (A) mRNA *in situ* hybridization. (B) GFP expression from an individual heterozygous for the cebpa^KI^ allele. For both methods, expression is strongest in the basal part of the inner dental epithelium.

**Figure S5:**
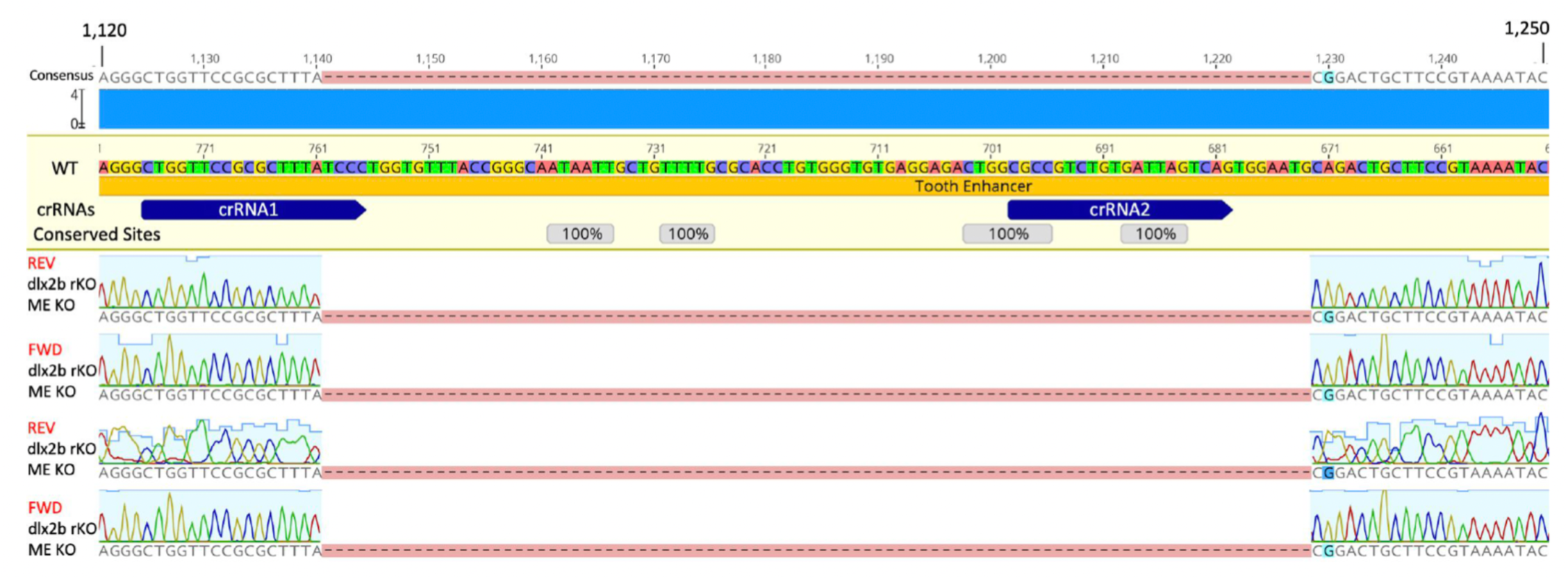
Sequence of the deletion in the dlx2b^KIΔMTE^ allele.

**Table S1.** Summary of reporter constructs used in this study. The dlx2b^MTE^ line is from construct #9. Mutation sequences are shown in Fig. S2.

| # | dlx2b<br>5'<br>end | dlx2b<br>3' end | Mutation | Type of<br>construct | Promote<br>r | Injected<br>molecule | Scored<br>as | Tooth GFP+ /<br>total<br>examined | Tooth activit<br>y |
| --- | --- | --- | --- | --- | --- | --- | --- | --- | --- |
| 1 | 1895 | 0 | none | Tol2kit<br>(Gateway) | dlx2b | plasmid | line | >100/>100 | + |
| 2 | 1555 | 0 | none | Tol2kit<br>(Gateway) | dlx2b | plasmid | F0 | 4/7 | + |
| 3 | 930 | 0 | none | Tol2kit<br>(Gateway) | dlx2b | plasmid | F0 | 3/4 | + |
| 4 | 856 | 0 | none | Tol2kit<br>(Gateway) | dlx2b | plasmid | F0 | 3/5 | + |
| 5 | 817 | 0 | none | Tol2kit<br>(Gateway) | dlx2b | plasmid | line | >100/>100 | + |
| 6 | 767 | 0 | none | Tol2kit<br>(Gateway) | dlx2b | plasmid | F0 | 0/19 | - |
| 7 | 622 | 0 | none | Tol2kit<br>(Gateway) | dlx2b | plasmid | F0 | 0/6 | - |
| 8 | 817 | 444 | none | Tol2kit<br>(Gateway) | β-actin | plasmid | line | >100/>100 | + |
| 9 | 817 | 604 | none | Tol2kit<br>(Gateway) | β-actin | plasmid | line | >100/>100 | + |
| 10 | 744 | 674 | none | Tol2kit<br>(Gateway) | β-actin | plasmid | line | 0/13 | - |
| 11 | 0 | 0 | none | Tol2kit<br>(Gateway) | β-actin | plasmid | line | 0/>100 | - |
| 12 | 817 | 604 | Pea3 | Tol2kit<br>(Gateway) | β-actin | plasmid | F0 | 12/35 | + |
| 13 | 817 | 604 | Cebp | NEBuilder | β-actin | amplicon | F0 | 3/5 | + |
| 14 | 817 | 604 | Dlx | NEBuilder | β-actin | amplicon | F0 | 3/7 | + |
| 15 | 817 | 604 | FoxA | NEBuilder | β-actin | amplicon | F0 | 6/14 | + |
| 16 | 817 | 604 | Ap1 | NEBuilder | β-actin | amplicon | F0 | 7/11 | + |
| 17 | 817 | 604 | HPM | NEBuilder | β-actin | amplicon | F0 | 12/14 | + |
| 18 | 817 | 604 | DFCA | NEBuilder | β-actin | amplicon | F0 | 0/30 | - |
| 19 | 817 | 604 | none | NEBuilder | hsp70 | plasmid | F0 | 5/6 | + |

